# Dopamine D3 Receptor Ligand Suppresses the Expression of Levodopa-Induced Dyskinesia in Nonhuman Primate Model of Parkinson’s Disease

**DOI:** 10.1101/2021.08.10.455884

**Authors:** Thomas Oh, Elyas S. Daadi, Jeffrey Kim, Etienne W. Daadi, Peng-Jen Chen, Gourav Roy-Choudhury, Jonathan Bohmann, Benjamin E. Blass, Marcel M. Daadi

## Abstract

Parkinson’s disease (PD) is a complex multisystem, chronic and so far incurable disease with significant unmet medical needs. The incidence of PD increases with aging and the expected burden will continue to escalate with our aging population. Since its discovery in the 1961 levodopa remains the gold standard pharmacotherapy for PD. However, the progressive nature of the neurodegenerative process in and beyond the nigrostriatal system causes a multitude of side effects, including levodopa-induced dyskinesia within 5 years of therapy. Attenuating dyskinesia has been a significant challenge in the clinical management of PD. We report on a small molecule that eliminates the expression of levodopa-induced dyskinesia and significantly improves PD-like symptoms. The lead compound PD13R we discovered is a dopamine D3 receptor partial agonist with high affinity and selectivity, orally active and with desirable drug-like properties. Future studies are aimed at developing this lead compound for treating PD patients with dyskinesia.

## INTRODUCTION

Parkinson’s disease (PD) is a relentlessly progressive neurodegenerative disorder impacting millions of people worldwide currently categorized as an eminent noninfectious pandemic [1]. In the US over a million people suffer from PD as an incurable disease and 60,000 new cases are diagnosed each year with an estimated cost of $27 billion per year [2]. Since the incidence of PD increases with aging, with an additional trend of increased incidence in men over the age of 70 [3], the expected burden will continue to escalate with our aging population.

PD pathology is associated primarily with the death of dopamine neurons in the substantia nigra and manifests with motor and non-motor dysfunctions including tremor, bradykinesia, rigidity, cognitive deficits and sleep disturbances [4–10].

These symptoms are managed by dopamine-replacement pharmacological treatments that enhance dopamine in the striatum with the dopamine precursor L-3,4-dihydroxy-phenylalanine (levodopa). Since it’s discovery in the 1961 [11, 12] (reviewed in [13]), levodopa remains the gold standard pharmacotherapy as it produces effective relief of the motor symptoms. However, the progressive nature of PD associated with the degenerative process within and beyond the nigrostriatal system causes a multitude of side effects within 5 years including the uncontrolled involuntary movements [14, 15] or levodopa-induced dyskinesia, the “wearing OFF” and the “ON/OFF” motor fluctuations. Over 80% of parkinsonian patients treated with levodopa develop levodopa-induced dyskinesia after five years [16, 17]. These debilitating side effects spurred the discovery of alternative dopamine-replacement pharmacological treatments dopamine receptor agonists and inhibitors of enzymes that degrade dopamine and the new anti-dyskinetic drug in use today, amantadine. Together, these alternative treatments help mitigate some of the side effects unfortunately each drug elicits a new range of additional side effects, which makes it challenging to manage the disease. Surgical treatments including deep brain stimulation (DBS) are alternatives that improve PD symptoms and dyskinesia. However, potential adverse events are associated with the surgical procedure and the device. The loss of DBS benefit parallels the progressive degenerative changes associated with PD.

In both rodent and nonhuman primate (NHP) models of PD with levodopa-induced dyskinesia, dopamine D3 receptor (D3R) expression is decreased during the development of PD-like symptoms and is abnormally up-regulated in the caudate and putamen after prolonged levodopa treatment and levodopa-induced dyskinesia expression [18, 19]. These data suggest the involvement of D3R in the pathogenesis of motor complications and levodopa-induced dyskinesia following levodopa pharmacotherapy.

Dopamine actions are mediated through five-member family of G-protein-coupled receptors (GPCRs) classified into two subtypes, D1-like and D2-like receptors based on sequence homology, G-protein coupling and pharmacology. D1R and D5R are coupled to the stimulatory D1-like G-protein family members (G_s/olf_), while D2R, D3R, and D4R are coupled to the inhibitory D2-like G-protein family members (G_i/o_). Although D3R has been implicated in PD and levodopa-induced dyskinesia, the high sequence homology between the D3R and D2R, including in the transmembrane region and the orthosteric binding site that binds dopamine, has made it challenging to translate promising leads to the clinic.

In this study we investigated a novel D3 receptor ligand, PD13R belonging to the arylpiperazine class of pharmacophore, to treat levodopa-induced dyskinesia in a NHP model of PD. PD13R has a high affinity and selectivity for D3R, and act as partial agonists via G-protein by inhibiting the adenylyl cyclase signaling pathway and decreasing cAMP production. The results demonstrated that PD13R inhibited the expression of levodopa-induced dyskinesia while improving motor functions and sleep efficiency in dyskinetic marmosets.

## MATERIALS AND METHODS

### Ethics Statement

The marmosets (Callithrix Jacchus) were from the Southwest Nonhuman Primate Research Center (SNPRC) colony. All the animals procedures were performed in strict accordance with the recommendations proposed in the Guide for the Care and Use of Laboratory Animals, National Research Council U. S. A. The protocols were approved by the Institutional Animal Care and Use Committee for Texas Biomedical Research Institute (approval no. 1461CJ, 1469CJ). All nonhuman primates held and used within the SNPRC program of care at the Texas Biomedical Research Institute are maintained under conditions that meet or exceed USDA Animal Welfare Regulations, OLAW standards, and National Institute of Health (NIH) guidelines as stated in the *Guide for the Care and Use of Laboratory Animals* (81h Edition, 2010), NAS-ILAR recommendations, and AAALAC accreditation standards for these species. Texas Biomed, including the SNPRC as a component of its overall program, is fully accredited by AAALAC International. The center promotes social housing caging, with structural complexities for environmental enrichment with the detailed observation of ongoing activities in the Animals. Animals are fed constant nutrition, complete life-cycle commercial monkey chows, supplemented daily with fruits and vegetables, and municipal drinking water is available at all times. All research activity has been conducted in accordance with the IACUC oversight process. The SNPRC employs a large number of full-time professional staff members to provide expertise in program administration, animal husbandry, clinical medicine, psychological well-being, facilities maintenance, animal records, and technical research support. All procedures were performed to minimize discomfort, distress or pain. Sedation and anesthetic agents are used to render the animal unconscious and therefore insensate to handling, discomfort, or pain. Analgesics are used to reduce any potential pain. All animals were enrolled in the environmental enrichment program. Enrichment provided to the animals consists of social contact, structural enrichment (e.g., perches, swings), manipulable enrichment (e.g., chew toys, balls), nutritional enrichment (e.g., fruit, grain), sensory enrichment (e.g., television, radio), and occupational enrichment (e.g., food puzzles). All enrichment provided is documented, and any deficiencies are addressed. Humane euthanasia of animals at the SNPRC is performed under the supervision of a staff veterinarian and in accordance with the professional principles and practices specified by the *American Veterinary Medical Association Guidelines for the Euthanasia of Animals: 2013 Edition.* Animals destined for euthanasia are injected intraperitoneally with sodium pentobarbital overdose (100 mg/Kg) followed by transcardial perfusion with phosphate buffered saline and 4% paraformaldehyde for tissue processing.

### Drug treatment

All the animals (n = 3) have been previously treated with MPTP as described in detail [20]. Briefly, MPTP (Sigma Aldrich) (dissolved in physiological saline) at a concentration of 2mg/ml was subcutaneously injected (2mg/Kg B.wt) for 5 consecutive days. After a wash out period of 72 hours from last MPTP injection the marmosets were returned to their home cages and monitored twice daily for rest of the study period.

For induction of dyskinesias, Levodopa (Sigma Aldrich) and carbidopa (Sigma Aldrich) at 1:1 were administered orally once daily for 5 days of a week. The drugs were mixed in either Ensure pudding, cottage cheese or marshmallows. The marmosets were started on a lower dose of 5mg/Kg B.wt during the first week and gradually increased to 10 and 15 mg/Kg B.wt over the course of next 2 weeks. Once the animals were acclimated to the drugs the dose was increased to 20 mg/Kg B.wt and continued for next 6 weeks.

We tested the anti-dyskinetic effects of 3 compounds: amantadine (Sigma Aldrich; 10mg/Kg B.wt), PD13R and SWR-3-73 both at 10mg/Kg (B.wt) synthesized as previously described [21, 22]. They were orally administered along with Levodopa and carbidopa (1:1, 20 mg/Kg B.wt) and the changes in dyskinesia symptoms and diurnal and nocturnal activities were analyzed using the dyskinesia disability score and the Actiwatch mini, respectively as described in the next section.

### Behavioral analysis

#### Parkinson’s Disease Rating Scale (PDRS)

The severity of symptoms in the marmosets was categorized using a validated parkinsonian rating scale for NHP [20]. The PDRS has been shown previously to correlate highly with striatal dopamine concentrations detected by postmortem immunohistopathology [20] and it is modeled on the Unified Parkinson’s Disease Rating Scale (UPDRS) [23] and the Hoehn & Yahr scale used clinically to categorize PD patients [24]. The PDRS was performed in daylight by video recording animals for 30 minutes. The evaluation was carried out biweekly before and after MPTP injections. The videos were scored by a blinded operator using PDRS, with a maximal disability score of 57 in the following manner: 0 = normal, 1 = mild, 2 = moderate, 3 = severe; rest tremor, action tremor, tremor of the head, tremor right arm, tremor left arm, freezing, locomotion, fine motor skills right hand, fine motor skills left hand, bradykinesia right arm, bradykinesia left arm, posture, hypokinesia, balance, posture, startle response, gross motor skills right hand and gross motor skills left hand, apathy (defined as a state of indifference), vocalization, drooling or frothing, tongue/face/lips. Independently of the PDRS, rigidity was assessed by evaluating the resistance to passive joint movements and the range of motion during reaching for food. Prior to MPTP lesion, we trained animals using their favorite food (i.e. marshmallows) as reward. The marmosets were trained and conditioned to perform the rewarding visually guided task of reaching and grabbing a marshmallow. The evaluation was performed before and after MPTP.

#### Dyskinesia Rating Scale

The severity of dyskinesia in the marmosets was categorized using a validated dyskinesia rating scale for NHPs [25–27]. Dyskinesia was analyzed from 5-minute videos of the monkeys recorded at every 30 minutes on Tuesday, Wednesday and Thursday of each week. The videos were recorded between 9:30 AM - 4:30 PM. Each day the compounds were administered at 11:30 AM. The severity of dyskinesias was scored for different segments of the animal’s body on a scale from 0 to 3, with 0 = absent, 1 = mild, 2 = moderate, 3 = severe. The body segments scored were: 1. Right upper limb dyskinesias (0-3); 2. Left upper limb dyskinesias (0-3); 3. Right lower limb dyskinesias (0-3); 4. left lower limb dyskinesias (0-3); 5. Trunk dyskinesias (0-3); 6, Head/facial dyskinesias (0-3). The severity of the rating was based on the frequency and amplitude of the abnormal movements. The total score is deduced from the six sub-scores to yield a dyskinesia score of 0 to 3: with 0 = absent, 1 = mild, manifesting by transient abnormal involuntary movements with choreic form, flicking of arms and limbs, increased running and hopping in the cages. 2 = moderate, characterized by frequent (over 50%) and intermittent uncontrolled irregular movements of limbs interfering with normal activity, while the animals are able to reach for treats. 3 = severe, characterized by a sustained dyskinesia manifesting with chorea and athetosis with sinuous writhing movements of the limbs, and dystonia with sustained extension of hind limbs and knees. The animals are unable to reach and grasp for treats (Video 1).

#### The object retrieval task with barrier detour (ORTBD)

The object retrieval task with a barrier detour is a reward based behavioral testing system that we previously described to evaluate motor and cognitive functions of NHP [20, 28]. Briefly, the task requires the test subject to retrieve a reward (marshmallow) fastened to a tray from the open side (bypassing the barrier) of a transparent box. For the current study, the testing apparatus was modified to fit the marmoset’s home cage and the animals were acclimatized to the apparatus prior to testing. Behavioral analysis was done for 3 consecutive days with 20 trials per day before and 1hr after administration of the test drugs. All parameters measured were previously described in detail [28]. During each trial the orientation of the open side of the box was randomly changed to either left or right of the animal or towards the opening of the cage. The entire process was recorded using a video camera and the recordings were then analyzed and scored by a blinded experimenter. During each trial, the following responses were scored (1) ability of the animal to reach the front, left, or right side of the box, scored under the term “reach act”; (2) hand of choice for the reach, either left or right, scored under the term “hand used”; (3) the outcome of the reach, either success or failure, scored under result section.

Using the above parameters, we were able to analyze the following additional variables: 1) Motor Problems: Reaching into the open side of the box but without retrieving the reward. 2) Initiation Time: Latency from the screen being raised to the subject touching the box or reward. 3) Execution: Retrieving the reward from the box on the first reach of the trial (indicates competence on the task). 4) Correct: Eventually retrieving the reward from the box on the trial (>1 reach on the trial to retrieve the reward because unlimited reaches per trial were allowed). 5) Reach number: Number of times the animal made an attempt and touched the box. 6) Hand preference: Hand (left or right) subject used for the first reach of the trial. 7) Hand bias: Total number of left and right hand reaches on each trial. 8) Awkward reach: Reaching with the hand farthest away from the box opening (either the left or right side). 9) Perseverative response: Repeating a reach to the side of the box that was previously open but then closed. 10) Barrier reach: Reaching and touching the closed side of the test box. The results from the data analysis were plotted using Graph pad prism statistical software.

#### Activity and sleep analysis

The diurnal and nocturnal behavior of the marmosets was monitored using the actiwatch mini (Cam*n*tech, UK) as previously described [20]. The device is an accelerometer that measures the intensity of the test subject’s omnidirectional movements in units or counts that are directly proportional to the animal’s activity. The device (2 cm in diameter) was placed on a collar around the neck of the marmoset. The animals were acclimatized to the collar in short sessions of 15 min followed by gradual increments of 30 min, 1hr, 3hrs, 6hrs and 12hrs. Once acclimatized, the actiwatch mini was attached to the collar and the activity-rest data was recorded for a period of 24 hours on 3 separate days of each treatments. The actiwatches were placed on animals at 8:30 AM and the devices were preset to start recording activity data from 9:00 AM. At 11:30 AM either the vehicle or the drugs were orally administered to the animals. The following day the actiwatch was removed after 9:00 AM and the data was transferred to a computer through an actiwatch reader using the actiwatch activity and sleep analysis-7 software (Cam*n*tech, UK). For sleep analysis, the period of sustained quiescence (marmosets sleep cycle) starting at 7:00 PM in the evening to early morning 6:30 AM (approximately 11 ½ hrs.) was analyzed using the sleep analysis-7 software to quantify the sleep quality and wakeful periods. The duration of sleep time was corrected for individual variations in the animal’s behavior to fall asleep at different time of the evening, thereby keeping the period of sleep time analyzed the same for all the animals. The analyzed data was then exported to excel and plotted using the Graph pad prism statistical software. To determine the change in activity after the administration of drugs actiwatch data in 10-minute bins were exported using the actiwatch activity and sleep analysis-7 software and plotted in Graph Pad Prism.

### Statistical analysis

Statistical analysis was done with Graph Pad Prism statistical software. Significance in differences between 2 groups was performed by applying Student’s t-test where appropriate. For comparison of multiple groups Two-Way ANOVA with Bonferroni post-hoc analysis was performed to identify the significant differences. A P-value of less than 0.05 was considered to be statistically significant.

## RESULTS

### Docking Simulation for PD13R reveals selectivity for D3R versus D2R

We have identified a potent dopamine D3 receptor (D3R) ligand, PD13R that is highly selective for the D3R over other dopamine receptors. We first performed molecular modeling studies to determine the binding region of PD13R and the D3R binding pocket. We used the Rhodium protein docking simulation program to implement a fully automated search over the agonist-bound D3R cryo-electron microscopy (cryo-EM) structure (PDB ID 7CMV) and predict the binding site. Rhodium’s unique docking pose analysis is based on changes in free energy during ligand-protein interaction, which is utilized in affinity optimization in ligand design [29, 30]. The model focuses on cavity-filling and on matching hydrogen-bond donors and acceptors at the ligand-protein interface, rather on optimizing their counts. The docking site selection is based on the long-range potential rules [31] and do not take in consideration analyst-chosen pockets. The optimal poses of PD13R in the agonist-bound cryo-EM structure revealed the structure of D3R bound to PD13R and localization of the amide group between TYR 365 and ASP 110 with polar contacts (Fig. 1A). The Asp-110 residue, located deep in the pocket represents a crucial pharmacological site of the PD13R binding at the orthosteric binding site (OBS) of D3R [32]. The lowest-energy pose with maximum cavity-filling is characterized by a short polar contact of the amide linker made with TYR 365 and by a longer contact made with SER 182 by the cyano group (Fig. 1B). Comparison with the off-target D2R cryo-EM structure (Fig. 1D) yields insights into selectivity of PD13R for D3R. In an overlay with the D2R structure (PDB 7JVR) the polar contacts to the key residues TYR 365 and SER 182 are lost. The TYR 365 became TYR 408 in D2R and reoriented completely away from the pocket leading to loss of the polar contact with the ligand. The lack of selectivity for D2R caused by the impairment of hydrogen bonding in the amide group of PD13R leads to a solvation penalty [30] not present in the on-target D3R. Thus selectivity is much higher towards D3R vs D2R because of the desolvation penalty for PD13R in D2R.

**Figure 1.**
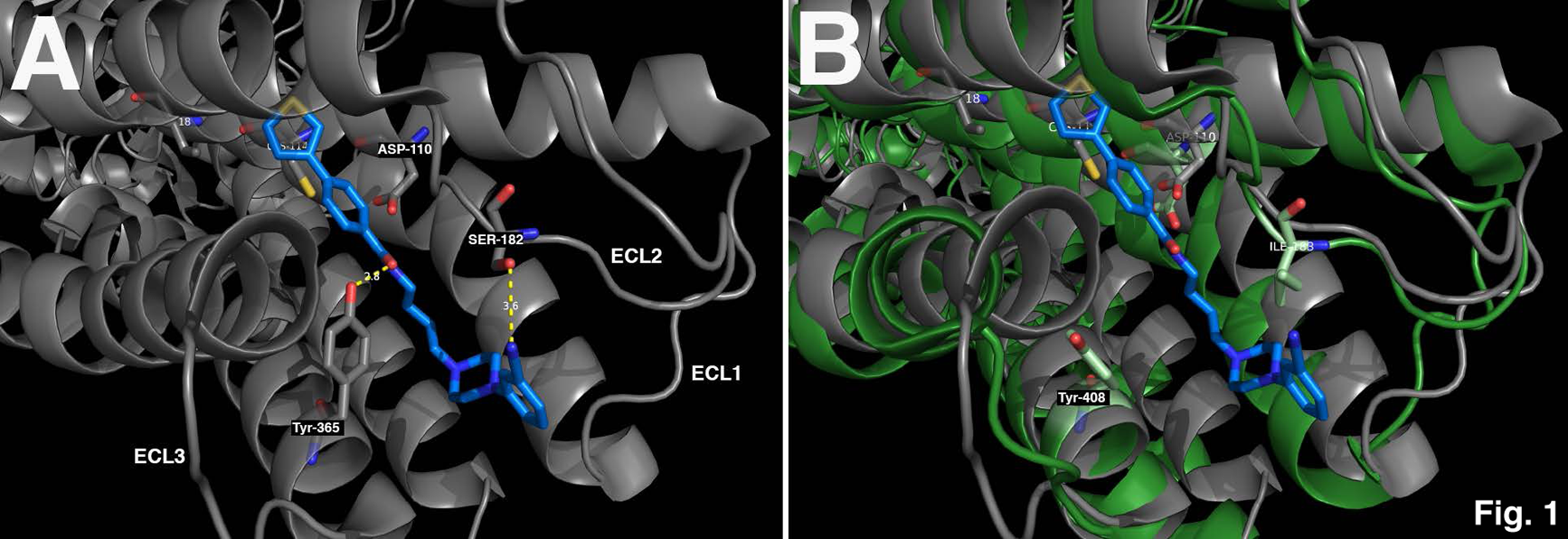
Docking Simulation for PD13R in D3R and D2R. **A**) Molecular recognition of PD13R (blue) in D3R cryo-EM structure showing interactions between the cyano (CN) group of the PD13R and residues TYR 365 and ASP 110 at the orthosteric binding site. There were polar contacts between the PD13R and the D3R including with the TYR 365 and ASP 110 residues. The pose shows short contact with TYR 365 and longer-range cyano group contact to SER 182 in flexible loop at pocket opening. **B** ) Structural superposition of PD13R-bound D2R (blue) demonstrating the difference in cavity filling between the 2 protein receptors D3R and D2R. D2R residues TYR 403 and ILE 283, representing S183I point mutation in D2R. Two potential points of polar contact with the ligands are lost.

### Pharmacology and Mechanism of action of PD13R

PD13R was tested in the in vitro competitive D3R and D2R radioligand binding assays and rat liver microsome (RLM) stability assay. We observed that the D3R tolerated better the electron donating and electron withdrawing substituents on the phenyl ring of the arylpiperazine as indicated by the potent and high sub-nanomolar affinity for D3R with Ki = 0.50 nM. The D3R over D2R (D3R Ki=19.8 nM, D2R Ki>17,000 nM, D2R/hyperD3R > 858).

Structure activity relationship (SAR) comparison between D3R versus D2R demonstrated that the incorporation of the cyano group in 3-position led to high affinity for the D3 receptor (Ki = 0.50 nM) and weaker affinity for the D2 receptor (D2 Ki = 743 nM) and a 1486 fold high selectivity for D3R over D2R. This data is consistent with our docking results demonstrating that the phenylpiperazine region is critical to the binding, while the high degree of selectivity for D3R observed was driven by the optimal cavity filling influenced by the length of the spacer and the phenyl thiophene region.

To determine the selectivity of PD13R toward other DA receptors we performed an in vitro screen for D1R, D4R, and D5R binding. The results showed a 1486 fold high selectivity for D3R over D2R, >20,000-fold selectivity for D3R over D1R or D5R and >1600-fold selectivity over D4R.

We then investigated the ability of the compound PD13R to couple G-proteins in the forskolin-dependent adenylyl cyclase inhibition assay and the β-arrestin binding assay. We used quinpirole and haloperidol as a full agonist and antagonist, respectively. Compounds were tested for efficacy in these assay at a dose equal to 10Å∼ their D3R Ki values in order to ensure > 90% receptor occupancy and the results were compared with the impact of the full agonist. There was no activity in the β-arrestin binding assay and partial agonism in the forskolin-dependent adenylyl cyclase inhibition assay producing 19.4% (mean Å} 5.4 S.E.M., n=3) of the activity of the maximum response observed with quinpirole.

### Development of levodopa-induced dyskinesia in Parkinsonian marmosets

We previously characterized the motor and non-motor Parkinson’s-like symptoms in the MPTP marmoset model [20] including postural and action tremors, altered range of motion during reaching, bradykinesia and problem solving during the skilled action of retrieving a reward. These symptoms were improved with levodopa treatment [20]. In this study we set to model levodopa-induced dyskinesia, a debilitating side effect that PD patients develop after long-term levodopa pharmacotherapy. Previous studies have demonstrated that the marmosets readily develop levodopa-induced dyskinesia after extended period of levodopa treatment [33, 34]. To induce dyskinesia we started the parkinsonian marmosets on a low dose of levodopa / Carbidopa (1:1, 5mg/Kg. B. Wt.) followed by medium dose (10 & 15 mg/Kg. B. Wt.). Once the animals acclimatized to the drugs, they were switched to high dose (20 mg/Kg. B. Wt.) and maintained for 6 weeks. To measure dyskinesia we used Dyskinesia disability rating scale that is scored for different segments of the animal’s body including face, trunk, arms and legs (see method section) and a wearable device, the actiwatch-mini [20] to monitor the animals’ activities while they are dyskinetic. During the low and medium dose of levodopa therapy the parkinsonian marmosets displayed a generalized increase in motility, absence of action tremors and increase in vocalization with no apparent symptoms of dyskinesias. In the first week of high dose levodopa treatment, the parkinsonian marmosets started displaying mild dyskinesia (Fig. 2) that manifested by transient abnormal movements, increased running and hopping in the cages. The majority of these abnormal involuntary movements were choreic with abnormal flicking of limbs and arms, which became visible approximately 30 minutes after dosing and lasted up to 180 minutes. In the following weeks however, the levodopa-induced dyskinesia gradually became more generalized with a steep increase in severity and total duration of symptoms (Fig. 2). The On-phase of levodopa lasted up to 4 hours after dosing with 3 well-delineated periods of dyskinesia, the onset period, the peak-dose period with maximum dyskinesia disability score and the end-of-dose period (Fig. 2). Within 30 minutes of dosing, the onset period, the marmosets started to manifest marked abnormal involuntary flowing movements of arms and limbs frequently tapping or touching cage walls or nest box. During the peak dose period the parkinsonian marmosets exhibited highest dyskinesia score and hyperactivity measured by actigraphy (Fig. 2A, B). The dyskinesia consisted of sustained hyperkinesia with stereotypic repetitive waving movements of the forearms and limbs [Video 1]. Chorea and athetosis (sinuous writhing movements of the limbs) were increasingly common at this stage. Dystonic movements were exhibited as sustained extension of hind limbs and knees.

**Figure 2.**
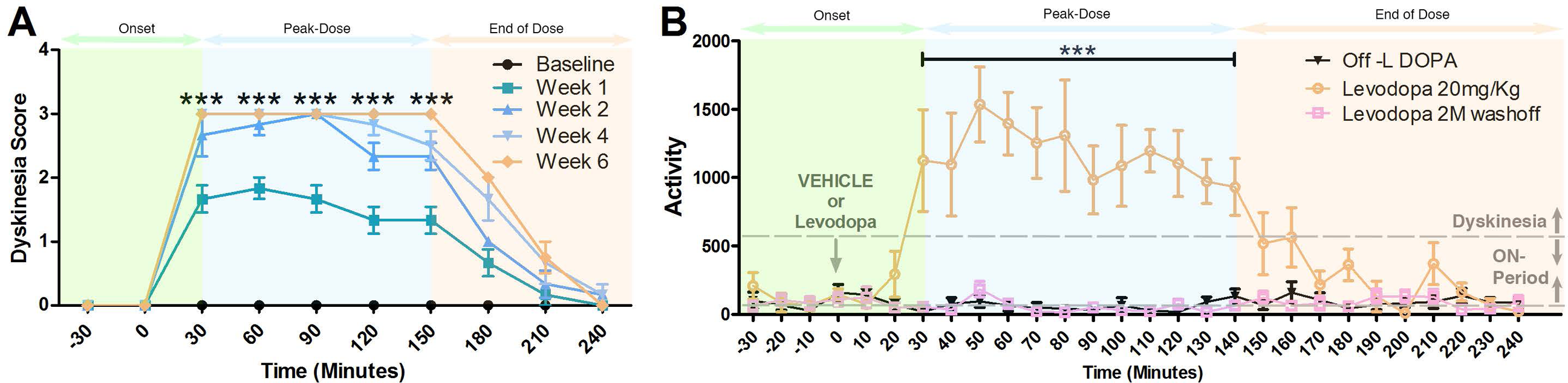
Parkinsonian marmosets Develop of levodopa-induced dyskinesia. To develop dyskinesia marmosets were treated with increasing dose of levodopa /Carbidopa (1:1) starting with doses of 5, 10 and 15 mg/Kg followed by a maximum dose of 20 mg/Kg, which was maintained for 6 weeks. A) Dyskinesia was measured weakly from week 1 of treatment to week 6 and analyzed as described in the Materials and Method section. B) Quantitative analysis of the abnormal activity of the animals registered by the actiwatch-mini during the period of dyskinesia induced in parkinsonan marmosets previously exposed to levodopa and acutely challenged with high dose 20 mg/kg of levodopa, compared to vehicle treated and 2 months (M) wash-off period. Arrow indicates time of the administration of vehicle or levodopa to the animals. Levodopa treatment resulted in a marked increase in motor activity of the parkinsonian marmosets reaching the ON-period and crosses the dyskinesia threshold (dotted line) within 30 min after injection. At this stage, the peak-dose dyskinesia and the ON response coincide. The animals return to the ON-period without dyskinesia approximately 145 minutes after Levodopa treatment and gradually enter the wearing-OFF period. Statistical analysis was performed using two-way ANOVA followed by Bonferroni post-hoc test analysis. ***P < 0.001. Error bars represent standard error.

### Treatment with the dopamine D3R ligand PD13R prevents the expression of levodopa-induced dyskinesia in the parkinsonian marmosets

We previously characterized the motor and non-motor Parkinson’s-like symptoms in the MPTP marmoset model [20] including postural and action tremors, altered range of motion during reaching, bradykinesia and problem solving during the skilled action of retrieving a reward. These symptoms were improved with levodopa treatment [20]. Marmosets were treated with dyskinesia-inducing high dose of levodopa (20 mg/kg) in combination with either vehicle or increasing doses of PD13R. Animal treated with levodopa combined with vehicle reached the peak-dose period of dyskinesias in 30 min after dosing and manifested by marked abnormal involuntary flowing movements of arms and limbs frequently tapping or touching cage walls or nest box and a highest dyskinesia disability score of 3 that lasted for 2 hours. The parkinsonian marmosets showed that the peak-dose dyskinesia and the ON response coincide following Levodopa treatment. This outcome of levodopa treatment is consistent with the outcome observed in patients with levodopa-induced dyskinesias [35]. When the parkinsonian marmosets were treated with levodopa (20 mg/kg) and PD13R at dose of 0.1 mg/kg no anti-dyskinetic effects were observed. While treatment with levodopa + PD13R at 1 mg/kg dose reduced the duration of the peak-dose period to 1.5 hours with a progressive decrease during the end-of-dose period in dyskinesia disability score (Fig. 3A). The combination of levodopa + PD13R at 5 mg/kg dose reduced the expression of levodopa-induced dyskinesia during the onset period, reaching peak-dose after 1 hour and reduced the peak levodopa effects to 1 hour before starting a progressive decrease (Fig. 3A). When used at a dose of 10 mg/kg in combination with levodopa, PDR13 drastically reduced the expression of dyskinesia by approximately 85% in the peak-dose period of levodopa-induced dyskinesia (Fig. 3A).

**Figure 3.**
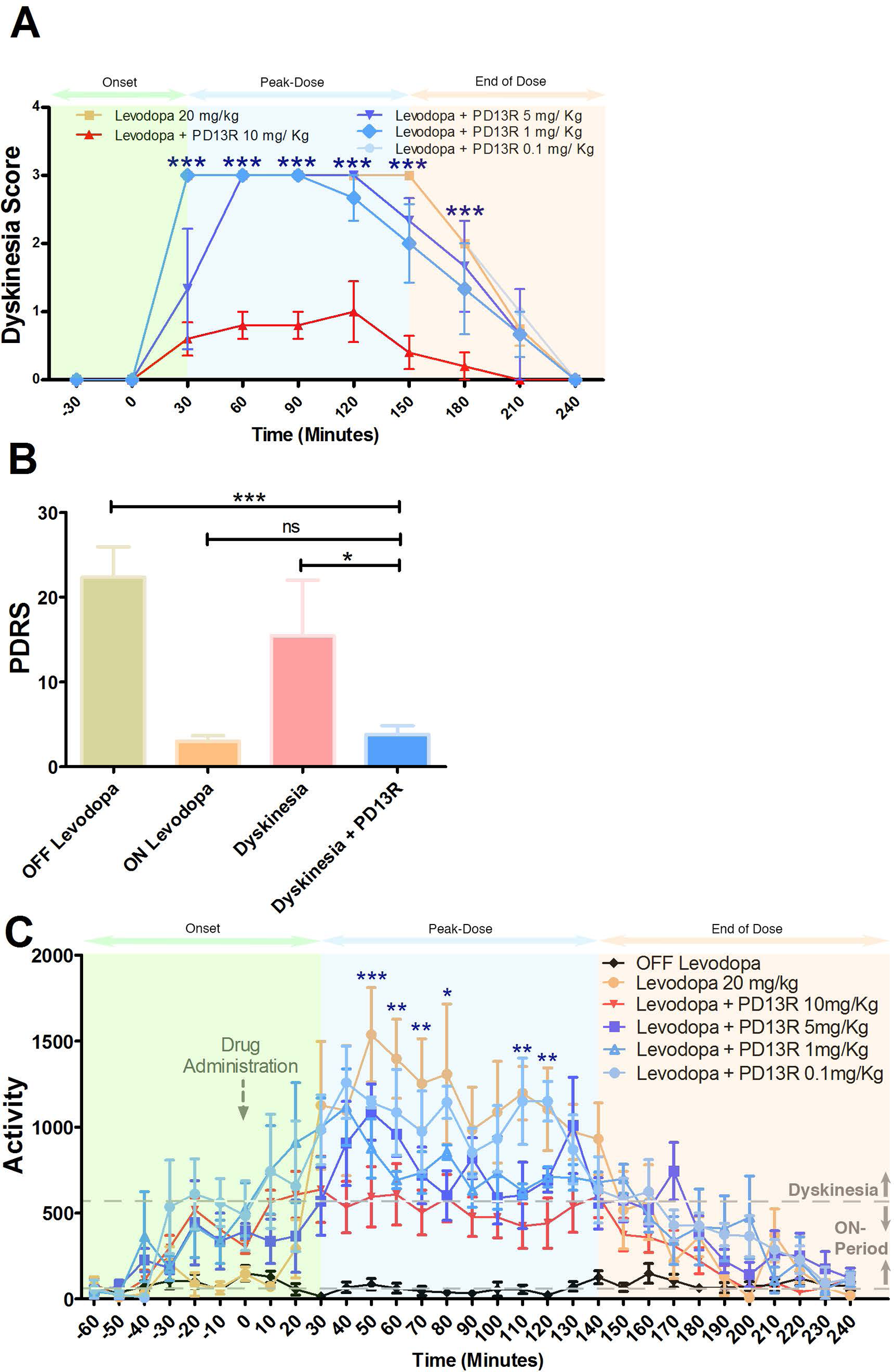
Treatment with the dopamine D3R ligand PD13R prevents the expression of levodopa-induced dyskinesia in the parkinsonian marmosets. A) Dose-range analysis of PD13R anti-dyskinetic effects. Dyskinesia was induced in parkinsonian marmosets previously exposed to levodopa by acute challenge with 20 mg/kg of levodopa combined with increasing doses of PD13R compound. The anti-dyskinetic effects of PD13R were dose-dependent with optimal outcome at 10 mg/Kg. B) PD13R eliminated the levodopa-induced dyskinesia at the optimal dose (10 mg/Kg) and improved the PDRS in dyskinetic parkinsonian marmosets. C) Quantitative analysis of the motor activity of the animals recorded by the actiwatch-mini during levodopa-induced dyskinesias in marmosets treated with 20 mg/kg of levodopa in the presence of increasing doses of PD13R. Arrow indicates time of drug administration. Levodopa treatment resulted in a marked hyperactivity that surpassed the therapeutic ON- period and crossed the dyskinesia threshold (dotted line) during the first 50 min after drug administration for all doses except the therapeutic dose 10 mg/Kg. At the later dose, motor activity remained within the therapeutic ON-period zone without dyskinesia. Statistical analysis was performed using two-way ANOVA followed by Bonferroni post-hoc test analysis. *P < 0.05, **P < 0.01, ***P < 0.001, ns: Not Significant. Error bars represent standard error.

### PD13R improved the PDRS and normalized daily activity of the parkinsonian marmosets with levodopa-induced dyskinesia

We then asked whether or not the anti-dyskinetic effects of PD13R led to the improvement of the Parkinson’s disease-like symptoms. We first used the PDRS to evaluate the parkinsonian syndrome during levodopa-induced dyskinesia. The parkinsonian marmosets displayed resting tremors, bradykinesia, hypokinesia and apathy with an average PDRS of 25 (Fig. 3B). High doses of Levodopa, induced dyskinesias and worsened the parkinsonian symptoms while physiological doses improved the PDRS (Fig. 3B). Importantly PD13R treatment eliminated the levodopa-induced dyskinesia and significantly improved the PDRS (Fig. 3B). To further confirm the anti-parkinsonian effects of PD13R, we used the actiwatch-mini and measured the animals’ activities during dyskinesia. The parkinsonian marmosets were treated with increasing doses of PD13R (0.1 to 10 mg/kg) in combination of dyskinesias-inducing high dose of levodopa (20 mg/kg). The actograms revealed a dose-dependent and significant reduction in the hyperactivity during the peak-dose period of levodopa-induced dyskinesia (Fig. 3C). PD13R improved the hyperactivity at 10 mg/kg and kept the motor activity within the ON-period with optimum anti-parkinsonian and anti-dyskinetic beneficial effects enabling the animal to move around freely with motor complications (Video 2).

### Comparative anti-dyskinetic effects of PD13R and amantadine

Amantadine is currently the main drug used in clinical practice under the name “GOCOVRI” or “ADS-5102” for treating levodopa-induced dyskinesias [36, 37]. We therefore set to investigate the anti-dyskinetic beneficial effects of PD13R compared to amantadine as a positive control and to a second D3R receptor partial agonist SWR-3-73 [22]. Parkinsonian marmosets were treated with dyskinesia-inducing high dose of levodopa (20 mg/kg) in combination with either amantadine (10 mg/kg) or PD13R (10 mg/kg). The data showed that both amantadine and PD13R significantly inhibited the expression of dyskinesias throughout the onset, peak-dose and end-of-dose periods of levodopa treatment (Fig. 4A).

**Figure 4.**
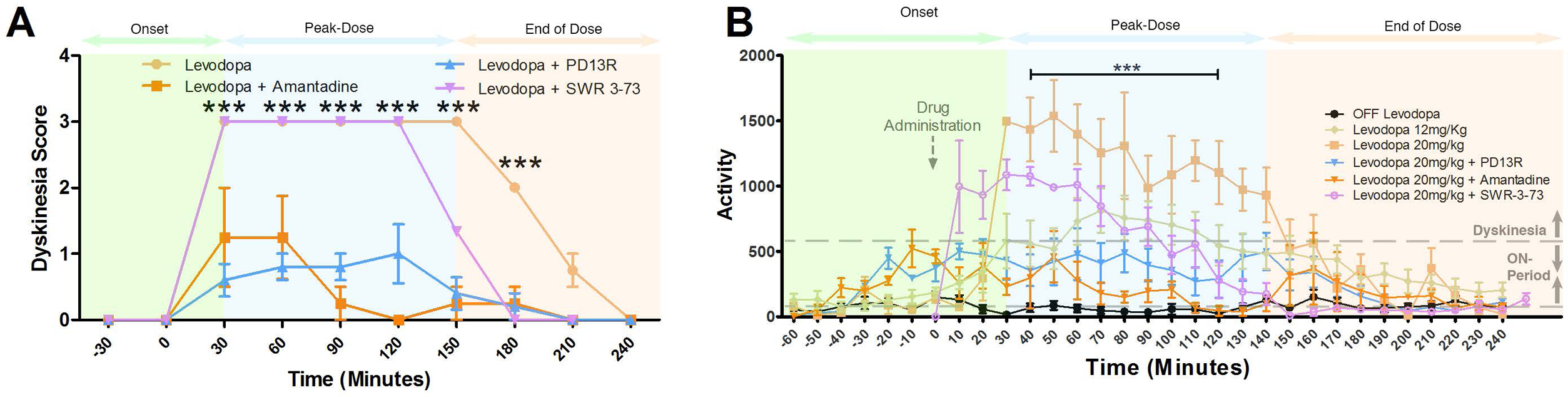
Anti-dyskinetic effects of PD13R compared to the NMDA glutamate receptor inhibitor amantadine. A) Dyskinesia was induced in parkinsonian marmosets previously exposed to levodopa by acute challenge with 20 mg/kg of levodopa combined with therapeutic doses (10 mg/Kg) of: 1) PD13R with arylpiperazine-based pharmacophore versus 2) SWR-3-73 with diazaspiro-based pharmacophore and 3) amantadine. C) Quantitative analysis of the time course motor activity of the animals recorded by the actiwatch-mini during levodopa-induced dyskinesias in marmosets treated with 20 mg/kg of levodopa in the presence of PD13R, SWR-3-73 and amantadine. Arrow indicates time of drug administration. At the therapeutic dose of 10 mg/Kg dose, PD13R and Amantadine inhibited the expression of dyskinesia compared to SWR-3-73 treatment. Both PD13R and Amantadine maintained the motor activity within the ON-period without dyskinesia. Statistical analysis was performed using two-way ANOVA followed by Bonferroni post-hoc test analysis. ***P < 0.001. Error bars represent standard error.

We next compared the anti-dyskinetic effects of PD13R and amantadine both at 10 mg/kg on the peak-dose levodopa-induced dykinesias using the actiwatch-mini (Fig. 4B). Both PD13R and amantadine inhibited the increased hyperactivity exhibited by the animals during the peak-dose period of levodopa-induced dyskinesias.

### Amantadine causes sleep disturbances in dyskinetic parkinsonian marmosets under levodopa treatment

The actiwatch enable us to non-invasively and empirically analyze the activity and the quality of sleep during nighttime in dyskinetic marmosets treated with levodopa in combination with increasing doses of PD13R (0.1 to 10 mg/kg) or with amantadine. Quantitative analysis of the actograms demonstrated significant increase in nocturnal activity, in time spent moving and wakefulness at nighttime in dyskinetic animals treated with amantadine compared to vehicle to PD13R- treated animals (Fig. 5). We then investigated the soundness of sleep by analyzing the sleep efficiency and actual awake time during the assumed sleep period. The results showed that sleep efficiency significantly decreased in dyskinetic parkisnonian marmoset treated with amantadine compared to vehicle-treated group and to dyskinetic animals treated with PD13R (Fig. 6). Furthermore, the actual awake time was significantly increased in dyskinetic animals treated with amantadine compared to PD13R-treated animals suggesting the marmosets woke up more often at night under amantadine treatment. Together these results suggested that dyskinetic parkinsonian marmosets treated with amantadine experienced abnormal irregular sleep and sleep disturbances while both D3R partial agonists, SWR373 and PD13R did not affect sleep.

**Figure 5.**
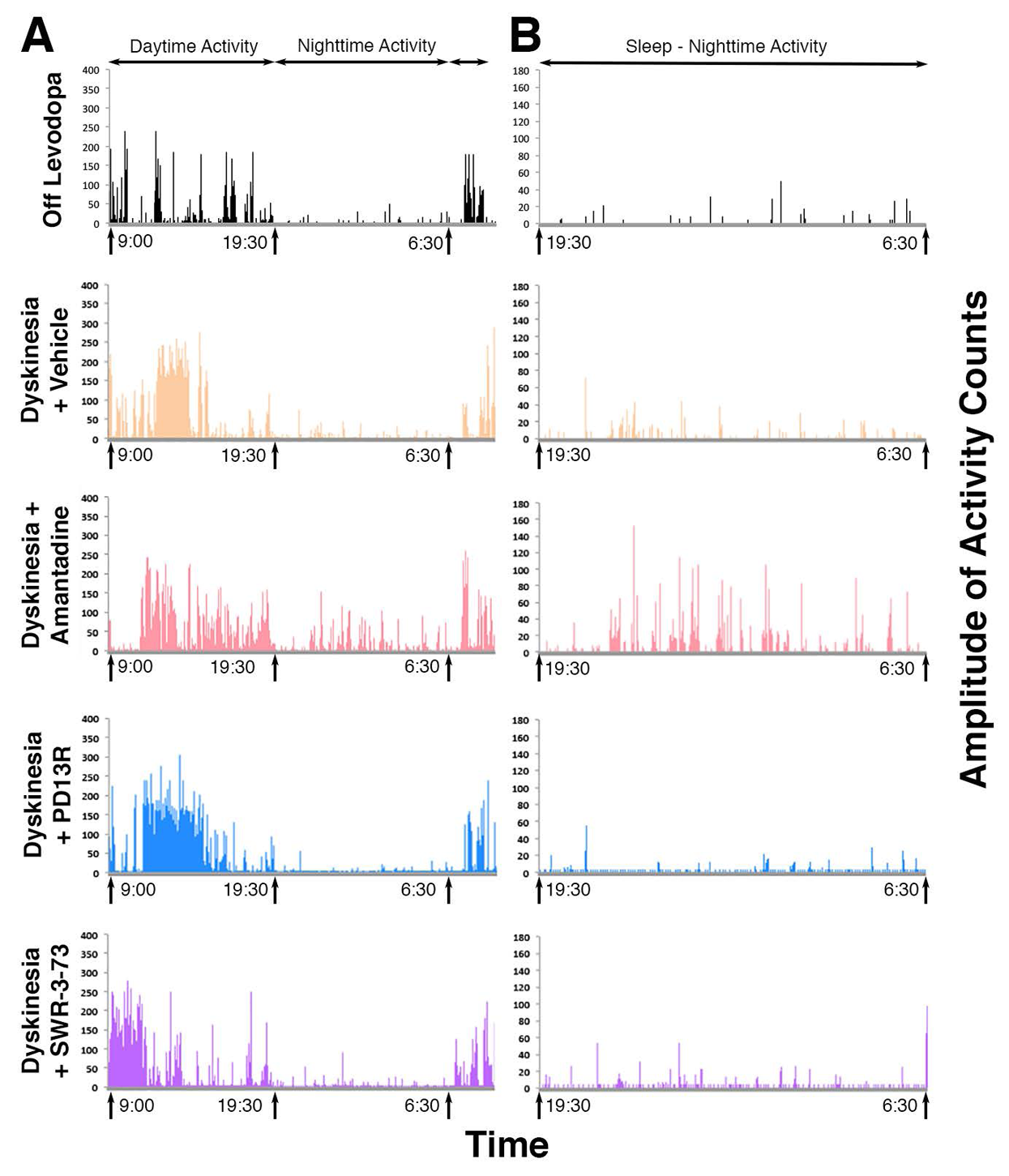
Amantadine causes sleep disturbances in dyskinetic parkinsonian marmosets under levodopa treatment. Representative actograms of parkinsonian marmosets throughout 24-hour daytime and nighttime period (9 am to 9 am next day) recording animal activities. (B) Representative actograms of the nocturnal activity (7 pm to 6:30 am). All actograms were recorded during the OFF period, and during levodopa-induced dyskinesias in marmosets acutely challenged with 20 mg/kg of levodopa in the presence of vehicle, PD13R, SWR-3-73 or amantadine. The data shows a clear abnormal increase in nocturnal activity in dyskinetic animals treated with amantadine while both PD13R and SWR-3-73 treated animals exhibited reduced baseline-level nocturnal activity.

**Figure 6:**
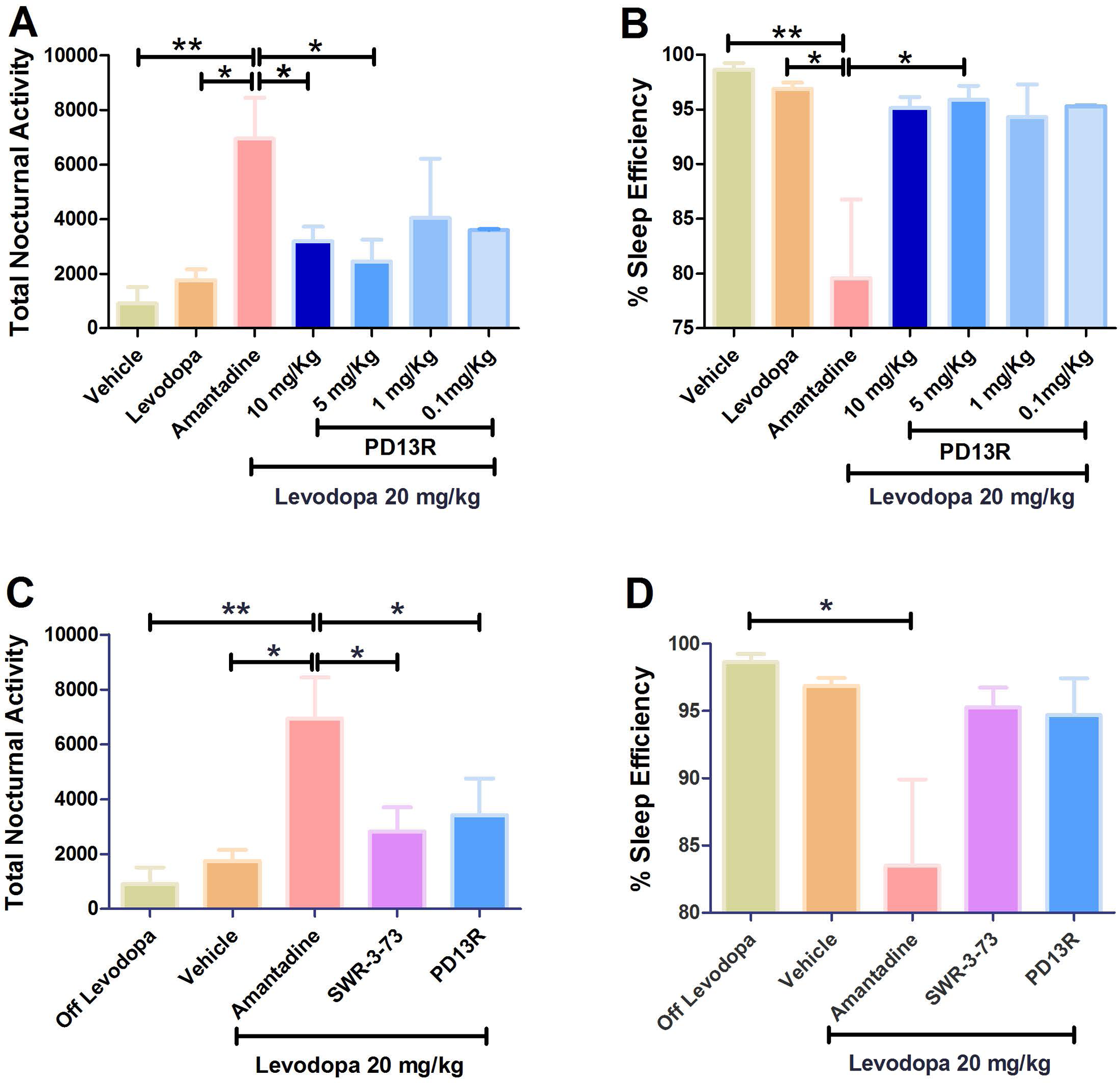
Amantadine worsens sleep quality in parkinsonian marmosets. A) & C) Quantitative analysis of sleep quality parameters showed that dyskinetic marmosets exhibited a significant increase in nocturnal activity under amantadine treatment, while neither PD13R under increasing doses (A) nor SWR-3-73 at 10 mg/Kg (C) modified the nocturnal activity of the animals. B) & D) the sleep efficiency also significantly dropped under amantadine treatment compared to PD13R administered at increasing doses (B) and to SWR-3-73 at 10 mg/Kg (C). Statistical analysis was performed using two-way ANOVA followed by Bonferroni post-hoc test analysis. *P < 0.05, **P < 0.01, ***P < 0.001. Error bars represent standard error.

#### DISCUSSIONS

We provide evidence that a novel D3 receptor partial agonist, PD13R with an arylpiperazine-based pharmacophore significantly suppresses the expression of levodopa-induced dyskinesia in a NHP model of PD. The anti-dyskinetic effects of PD13R didn’t affect the anti-parkinsonian benefits of levodopa while improving sleep efficiency compared to amantadine. PD13R demonstrated desirable CNS drug-like features with high affinity for the D3R (Ki = 0.50 nM), high selectivity for the D3R over the D2R (1486-fold), >20,000-fold selectivity for D3R over D1R or D5R and >1600-fold selectivity for D3R over D4R. PD13R exhibited low efficacy in the forskolin-dependent adenylyl cyclase inhibition assay (19.4% maximum activity) and a prolonged half-life (>60 m in human and rat liver microsome assays) of over 10 hrs in the in vivo PK study. Together these data demonstrated that the lead compound PD13R is efficacious in treating levodopa-induced dyskinesias, orally active with desirable drug-like properties, including potency, selectivity and bioavailability.

Levodopa-induced dyskinesia develops in up 95% of patients after 15 years of treatment [17, 38, 39] yet it remains the gold standard pharmacotherapy for PD. Thus, attenuating dyskinesia has been a significant challenge in the clinical management of Parkinson’s disease. The attempt to attenuate dyskinesia by reducing or adjusting the dose of levodopa and implementing other adjunctive dopamine-replacement drugs has produced inconsistent outcomes. Here we report a small molecule that eliminates the expression of levodopa-induced dyskinesia without affecting the anti-parkinsonian effects of Levodopa. The lead compound PD13R we discovered is a D3R receptor partial agonist demonstrated in the forskolin-dependent adenylyl cyclase inhibition assay producing 19.4% of the activity of the maximum response observed with quinpirole.

D3R plays a critical role in central nervous system health and disease. Although D3R involvement in drug abuse and addiction, reward, cognition, schizophrenia, impulse control disorders and Parkinson’s disease is widely understood, it has also been implicated in a variety of other functions including neuroinflammation, neurogenesis, protein aggregation and neurotrophic factor secretion (Reviewed in [40, 41]. The expression of D3R in Parkinson’s disease animal models and patients mirrors the dynamic state of the disease impacted by the developmental timeline of the PD-like symptoms, by the evolution of the neurodegenerative process and by therapeutic interventions such as the levodopa treatment. Previous reports have also suggested that D3R expression is species specific, in rat models the D3R expression is limited to dopaminergic areas known to be associated with cognitive and emotional functions [18, 42–45], while in the NHP models, the D3R expression closely resembles the human pattern with a widespread expression in motor and associative structures of the basal ganglia and in limbic system [19, 46–52]. In all these species however, D3R is down regulated in the basal ganglia after the stabilization of PD-like symptoms [19, 47, 53–55] and D3R increases in expression and binding in caudate, putamen and globus pallidus after levodopa treatment and development of levodopa-induced dyskinesia [18, 19, 47, 56, 57]. These data suggest that the D3R is involved in the genesis and expression of levodopa-induced dyskinesia and may represent a viable therapeutic target for its treatment.

In the CNS disorders, improving the D3 versus D2 compound selectivity has been the most challenging aspect in developing selective D3R ligands. The sequence similarity between D3R and D2R is 90% while sequence identities in relevant interactions sites with ligands such as the transmembrane region are highest for D2 versus D3 at 79% [58]. Our docking simulation analysis for PD13R revealed high selectivity for D3R versus D2R. The overlay of D3R and D2R crystal structures docking pose of the PD13R revealed a loss of the polar contact at the key residues TYR 365 and SER 182, the TYR 365 became TYR 408 and reoriented completely away from the pocket leading to loss of the polar contact with the ligand. Furthermore, the bottom part of the binding pocket is unoccupied in the D2R. This un-filled portion of the pocket is important because cavity-filling connects selectivity to affinity [29], and the unfilled void would decrease selectivity and binding affinity to D2R. The optimal cavity filling of PD13R for D3R vs D2R could be mediated by the spacer linking the arylpiperazine to the arylamid, enabling a full cavity filling and improved selectivity and affinity to D3R. These finding are consistent with previous SAR studies demonstrated that the length of the spacer impacts the affinity and selectivity to the D3R [59, 60]. Decreasing of the spacer length from butylene to propylene or ethylene markedly reduces affinity to D3R, similar results were obtained when the spacer is increased from butylene to pentamethylene (Reviewed in [60]). Interestingly, PD13R docked in an inverted conformation in the anti-dyskinetic agonist-bound cryo-EM structure of the D3R (7CMV) compared to the antipsychotic antagonist-bound X-Ray structure of the receptor (3PBL) [21]. The difference between the ’reverse’ and ’forward’ conformers by docking could be sensitive to conformer sampling. Further simulation studies are needed to correlate docking position to the potency of the compound.

Our data suggest that arylpiperazine is critical for the anti-dyskinetaic beneficial effects. Substitution in the 4-position of the phenyl ring of the arylpiperazine lead to losses in binding to D2R and a 1486 fold high selectivity for D3R over D2R. Together these data suggest that arylpiperazine, along with the spacer and the phenyl thiophene drives the optimal activity and cavity filling of D3R. These findings are consistent with the structure activity relationships for D3R antagonists in substance abuse concluding that in order to obtain high affinity for D3R (<10 nm), high selectivity over D2R (> 100 fold) the arylpiperazine-based scaffolds are the most promising [59, 61]. Specifically, the structural features for potent and selective D3 antagonists include 1) an extended aryl amide, 2) piperazine containing an aromatic or heteroaromatic moiety in position 4 and 3) 4-atom (n-butyl) linking chain between the two functional groups. Although the arylpiperazine element is the primary pharmacophore, the selectivity for D3R also depends on the length of the linker [60–63]. The arylpiperazine phamacophore was mostly described as antagonist, which has created confusion in the Parkinson’s disease field, as it is also a common structure of partial D3R agonist [61]. Our study is the first to report the robust efficacy of this class of compounds in eliminating levodopa-induced dyskinesias.

Our lead compound PD13R, a partial D3R agonist, exerts approximately 19% adenylyl cyclase inhibition of a full agonist. Importantly PD13R did not affect the therapeutic benefits of levodopa, manifested by the improvement in the Parkinson’s disease rating score. The PD-like symptoms evaluated included tremors, bradykinesia, and posture and general activity measured by an accelerometer device, the Actiwatch mini. In addition we measured motor and cognitive functions using the object retrieval task with barrier detour [20]. As demonstrated in the video we observed a clear improvement in all PD-like symptoms and movement coordination in animals treated with PD13R compared to control.

The anti-dyskinetic effects of PD13R were similar to those observed with amantadine treatment, the currently prescribed drug for dyskinesia that we used as positive control. Based on several clinical studies [64–66], amantadine, which is an NMDA glutamate receptor inhibitor, was designated as efficacious for the treatment of dyskinesia by the Movement Disorder Society Evidence Based Medical Review [67]. Interestingly we detected different effects on sleep patterns between amantadine and PD13R. The PD13R did not affect sleep efficiency, while animals treated with amantadine exhibited a significant increase in nocturnal activity and decrease in sleep efficiency. Previous reports has highlighted the potential side effects of amantadine with the most common include cardiovascular dysfunction, myoclonus, orthostatic hypotension, peripheral edema, urinary tract infection, nervousness, insomnia, anxiety, hallucinations, delirium, confusion, nausea and constipation [68, 69]. Thus the search for better anti-dyskinetic agents remains a critical medical need. This novel D3R partial agonist, PD13R is a promising lead for the treatment of dyskinesia. It has demonstrated a high sub-nanomolar affinity for the D3R (Ki = 0.50 nM), 1486-fold higher selectivity for the D3R over the D2R, 20,000-fold higher selectivity for D3R over D1R or D5R and over 1600-fold selectivity for D3R over D4R. PD13R has a prolonged half-life of over 10 hours in vivo, and a partial agonistic activity (19.4% maximum activity) in the forskolin-dependent adenylyl cyclase inhibition assay.

Further studies are needed to better understand the anti-dyskinetic and anti- parkinsonian effects of the arylpiperazine versus other pharmacophores. Our future studies are aimed at optimizing and developing the lead compound PD13R for treating patients with Parkinson’s disease.

## Acknowledgements

The authors wish to thank Dr. Robert H. Mach and Dr. Robert R. Luedtke for providing comments on the manuscript and also thank Dr. Mach for providing the D3 agonist compound. This work was supported by the Worth Family Fund, The Perry and Ruby Stevens Charitable Foundation, The Robert J. Jr. and Helen C. Kleberg Foundation, The Marmion Family Fund, The William and Ella Owens Medical Research Foundation, the National Institute on Aging R56 AG059284 and the NIH Primate Center Base grant (Office of Research Infrastructure Programs/OD P51 OD011133).

## Author contributions

T.O., E.S.D., J.K., E.W.D., P.J.C., S.W.R., G.C., J.B., B.E.B., M.M.D. collected data; performed the data analysis; provided comments on the manuscript, contributed reagents, M.M.D. wrote manuscript. All the authors read and approved the final manuscript.

## Video Legend

**Video 1**: The video recording show typical dyskinetic marmoset during the peak- dose period displaying athetosis and choreic dyskinesia with sinuous writhing movements of the limbs. There is a sustained hyperkinesia with stereotypic repetitive waving movements of the forearms and limbs.

**Video 2**: The video recording shows a vehicle-treated dyskinetic marmoset with motor problems then the same animal treated with PD13R. The marmoset became freely moving during the peak dose period showing suppression of dyskinesia after treatment with PD13R compound.

## References

1. Dorsey ER, Bloem BR. The Parkinson Pandemic-A Call to Action. JAMA Neurol. 2018;75(1):9–10. Epub 2017/11/14. doi: 10.1001/jamaneurol.2017.3299. PubMed PMID: 29131880.

2. Obeso JA, Rodriguez-Oroz MC, Rodriguez M, Lanciego JL, Artieda J, Gonzalo N, et al. Pathophysiology of the basal ganglia in Parkinson’s disease. Trends Neurosci. 2000;23(10 Suppl):S8-19. PubMed PMID: 11052215.

3. Savica R, Grossardt BR, Bower JH, Ahlskog JE, Rocca WA. Time Trends in the Incidence of Parkinson Disease. JAMA Neurol. 2016;73(8):981–9. doi: 10.1001/jamaneurol.2016.0947. PubMed PMID: 27323276; PubMed Central PMCID: PMCPMC5004732.

4. Espay AJ, Brundin P, Lang AE. Precision medicine for disease modification in Parkinson disease. Nat Rev Neurol. 2017;13(2):119–26. Epub 2017/01/21. doi: 10.1038/nrneurol.2016.196. PubMed PMID: 28106064.

5. Chaudhuri KR, Schapira AH. Non-motor symptoms of Parkinson’s disease: dopaminergic pathophysiology and treatment. Lancet Neurol. 2009;8(5):464–74. Epub 2009/04/21. doi: 10.1016/s1474-4422(09)70068-7. PubMed PMID: 19375664.

6. Jellinger KA. Neuropathology of sporadic Parkinson’s disease: evaluation and changes of concepts. Mov Disord. 2012;27(1):8–30. Epub 2011/11/15. doi: 10.1002/mds.23795. PubMed PMID: 22081500.

7. Braak H, Del Tredici K. Neuropathological Staging of Brain Pathology in Sporadic Parkinson’s disease: Separating the Wheat from the Chaff. J Parkinsons Dis. 2017;7(s1):S73-s87. Epub 2017/03/12. doi: 10.3233/jpd-179001. PubMed PMID: 28282810; PubMed Central PMCID: PMCPMC5345633.

8. Jellinger KA. Neuropathology of Nonmotor Symptoms of Parkinson’s Disease. Int Rev Neurobiol. 2017;133:13–62. Epub 2017/08/15. doi: 10.1016/bs.irn.2017.05.005. PubMed PMID: 28802920.

9. Lang AE, Lozano AM. Parkinson’s Disease. New England Journal of Medicine. 1998;339(15):1044–53. doi: 10.1056/NEJM199810083391506.

10. Braak H, Braak E, Yilmazer D, Schultz C, de Vos RA, Jansen EN. Nigral and extranigral pathology in Parkinson’s disease. J Neural Transm Suppl. 1995;46:15–31. Epub 1995/01/01. PubMed PMID: 8821039.

11. Birkmayer W, Hornykiewicz O. [The L-3,4-dioxyphenylalanine (DOPA)- effect in Parkinson-akinesia]. Wien Klin Wochenschr. 1961;73:787–8. Epub 1961/11/10. PubMed PMID: 13869404.

12. Cotzias GC, Van Woert MH, Schiffer LM. Aromatic amino acids and modification of parkinsonism. N Engl J Med. 1967;276(7):374–9. Epub 1967/02/16. doi: 10.1056/nejm196702162760703. PubMed PMID: 5334614.

13. Jankovic J. Parkinson’s disease: clinical features and diagnosis. J Neurol Neurosurg Psychiatry. 2008;79(4):368–76. Epub 2008/03/18. doi: 10.1136/jnnp.2007.131045. PubMed PMID: 18344392.

14. Cotzias GC, Papavasiliou PS, Gellene R. Experimental treatment of parkinsonism with L-Dopa. Neurology. 1968;18(3):276–7. Epub 1968/03/01. PubMed PMID: 4869717.

15. Cotzias GC, Papavasiliou PS, Gellene R. Modification of Parkinsonism-- chronic treatment with L-dopa. N Engl J Med. 1969;280(7):337–45. PubMed PMID: 4178641.

16. Jankovic J. Motor fluctuations and dyskinesias in Parkinson’s disease: clinical manifestations. Mov Disord. 2005;20 Suppl 11:S11–6. Epub 2005/04/12. doi: 10.1002/mds.20458. PubMed PMID: 15822109.

17. Ahlskog JE, Muenter MD. Frequency of levodopa-related dyskinesias and motor fluctuations as estimated from the cumulative literature. Mov Disord. 2001;16(3):448–58. Epub 2001/06/08. doi: 10.1002/mds.1090. PubMed PMID: 11391738.

18. Bordet R, Ridray S, Carboni S, Diaz J, Sokoloff P, Schwartz JC. Induction of dopamine D3 receptor expression as a mechanism of behavioral sensitization to levodopa. Proc Natl Acad Sci U S A. 1997;94(7):3363–7. Epub 1997/04/01. doi: 10.1073/pnas.94.7.3363. PubMed PMID: 9096399; PubMed Central PMCID: PMCPMC20375.

19. Bezard E, Ferry S, Mach U, Stark H, Leriche L, Boraud T, et al. Attenuation of levodopa-induced dyskinesia by normalizing dopamine D3 receptor function. Nat Med. 2003;9(6):762–7. Epub 2003/05/13. doi: 10.1038/nm875. PubMed PMID: 12740572.

20. Choudhury GR, Daadi MM. Charting the onset of Parkinson-like motor and non-motor symptoms in nonhuman primate model of Parkinson’s disease. PLoS One. 2018;13(8):e0202770. Epub 2018/08/24. doi: 10.1371/journal.pone.0202770. PubMed PMID: 30227600; PubMed Central PMCID: PMCPMC6107255 alter our adherence to PLOS ONE policies on sharing data and materials.

21. Chen PJ, Taylor M, Griffin SA, Amani A, Hayatshahi H, Korzekwa K, et al. Design, synthesis, and evaluation of N-(4-(4-phenyl piperazin-1-yl)butyl)-4- (thiophen-3-yl)benzamides as selective dopamine D(3) receptor ligands. Bioorganic & medicinal chemistry letters. 2019;29(18):2690–4. Epub 2019/08/08. doi: 10.1016/j.bmcl.2019.07.020. PubMed PMID: 31387791.

22. Reilly SW, Griffin S, Taylor M, Sahlholm K, Weng CC, Xu K, et al. Highly Selective Dopamine D(3) Receptor Antagonists with Arylated Diazaspiro Alkane Cores. J Med Chem. 2017;60(23):9905–10. Epub 2017/11/11. doi: 10.1021/acs.jmedchem.7b01248. PubMed PMID: 29125762; PubMed Central PMCID: PMCPMC5767125.

23. Fahn S, Elton R, Committee UD. Unified Parkinson’s disease rating scale, Fahn S., Marsden CD, Calne DB, Goldstein M., Recent developments in Parkinson’s disease vol. 2, 1987, 153–164. Macmillan Health Care Information, Florham Park, NJ.

24. Hoehn MM, Yahr MD. Parkinsonism: onset, progression and mortality. Neurology. 1967;17(5):427–42. Epub 1967/05/01. PubMed PMID: 6067254.

25. Bankiewicz KS, Daadi M, Pivirotto P, Bringas J, Sanftner L, Cunningham J, et al. Focal striatal dopamine may potentiate dyskinesias in parkinsonian monkeys. Experimental Neurology. 2006;197(2):363–72. PubMed PMID: WOS:000235131000012.

26. Quik M, Mallela A, Ly J, Zhang D. Nicotine reduces established levodopa-induced dyskinesias in a monkey model of Parkinson’s disease. Mov Disord. 2013;28(10):1398–406. doi: 10.1002/mds.25594. PubMed PMID: 23836409; PubMed Central PMCID: PMCPMC3787977.

27. Fox SH, Johnston TH, Li Q, Brotchie J, Bezard E. A critique of available scales and presentation of the Non-Human Primate Dyskinesia Rating Scale. Mov Disord. 2012;27(11):1373–8. Epub 2012/09/15. doi: 10.1002/mds.25133. PubMed PMID: 22976821.

28. McEntire CR, Choudhury GR, Torres A, Steinberg GK, Redmond DE, Jr., Daadi MM. Impaired Arm Function and Finger Dexterity in a Nonhuman Primate Model of Stroke: Motor and Cognitive Assessments. Stroke. 2016;47(4):1109–16. Epub 2016/03/10. doi: 10.1161/strokeaha.115.012506. PubMed PMID: 26956259.

29. Olsson TS, Williams MA, Pitt WR, Ladbury JE. The thermodynamics of protein-ligand interaction and solvation: insights for ligand design. J Mol Biol. 2008;384(4):1002–17. Epub 2008/10/22. doi: 10.1016/j.jmb.2008.09.073. PubMed PMID: 18930735.

30. Ladbury JE, Klebe G, Freire E. Adding calorimetric data to decision making in lead discovery: a hot tip. Nat Rev Drug Discov. 2010;9(1):23–7. Epub 2009/12/05. doi: 10.1038/nrd3054. PubMed PMID: 19960014.

31. Vakser IA. Long-distance potentials: an approach to the multiple-minima problem in ligand-receptor interaction. Protein engineering. 1996;9(1):37–41. Epub 1996/01/01. doi: 10.1093/protein/9.1.37. PubMed PMID: 9053900.

32. Chien EY, Liu W, Zhao Q, Katritch V, Han GW, Hanson MA, et al. Structure of the human dopamine D3 receptor in complex with a D2/D3 selective antagonist. Science. 2010;330(6007):1091-5. Epub 2010/11/26. doi: 10.1126/science.1197410. PubMed PMID: 21097933; PubMed Central PMCID: PMCPMC3058422.

33. Pearce RK, Jackson M, Smith L, Jenner P, Marsden CD. Chronic L-DOPA administration induces dyskinesias in the 1-methyl-4- phenyl-1,2,3,6- tetrahydropyridine-treated common marmoset (Callithrix Jacchus). Mov Disord. 1995;10(6):731–40. Epub 1995/11/01. doi: 10.1002/mds.870100606. PubMed PMID: 8749992.

34. Jenner P. Factors influencing the onset and persistence of dyskinesia in MPTP-treated primates. Ann Neurol. 2000;47(4 Suppl 1):S90–9; discussion S9-104. PubMed PMID: 10762136.

35. Encarnacion EV, Hauser RA. Levodopa-induced dyskinesias in Parkinson’s disease: etiology, impact on quality of life, and treatments. Eur Neurol. 2008;60(2):57–66. Epub 2008/05/16. doi: 10.1159/000131893. PubMed PMID: 18480609.

36. Shannon KM, Goetz CG, Carroll VS, Tanner CM, Klawans HL. Amantadine and motor fluctuations in chronic Parkinson’s disease. Clin Neuropharmacol. 1987;10(6):522–6. Epub 1987/12/01. doi: 10.1097/00002826- 198712000-00003. PubMed PMID: 3427558.

37. Verhagen Metman L, Del Dotto P, van den Munckhof P, Fang J, Mouradian MM, Chase TN. Amantadine as treatment for dyskinesias and motor fluctuations in Parkinson’s disease. Neurology. 1998;50(5):1323–6. Epub 1998/05/22. doi: 10.1212/wnl.50.5.1323. PubMed PMID: 9595981.

38. Hely MA, Morris JG, Reid WG, Trafficante R. Sydney Multicenter Study of Parkinson’s disease: non-L-dopa-responsive problems dominate at 15 years. Mov Disord. 2005;20(2):190–9. Epub 2004/11/20. doi: 10.1002/mds.20324. PubMed PMID: 15551331.

39. Olanow CW, Watts RL, Koller WC. An algorithm (decision tree) for the management of Parkinson’s disease (2001): treatment guidelines. Neurology. 2001;56(11 Suppl 5):S1-s88. Epub 2001/06/13. doi: 10.1212/wnl.56.suppl_5.s1. PubMed PMID: 11402154.

40. Yang P, Perlmutter JS, Benzinger TLS, Morris JC, Xu J. Dopamine D3 receptor: A neglected participant in Parkinson Disease pathogenesis and treatment? Ageing Res Rev. 2020;57:100994. Epub 2019/11/26. doi: 10.1016/j.arr.2019.100994. PubMed PMID: 31765822; PubMed Central PMCID: PMCPMC6939386.

41. Sokoloff P, Le Foll B. The dopamine D3 receptor, a quarter century later. Eur J Neurosci. 2017;45(1):2–19. Epub 2016/09/08. doi: 10.1111/ejn.13390. PubMed PMID: 27600596.

42. Lévesque D, Diaz J, Pilon C, Martres MP, Giros B, Souil E, et al. Identification, characterization, and localization of the dopamine D3 receptor in rat brain using 7-[3H]hydroxy-N,N-di-n-propyl-2-aminotetralin. Proc Natl Acad Sci U S A. 1992;89(17):8155-9. Epub 1992/09/01. doi: 10.1073/pnas.89.17.8155. PubMed PMID: 1518841; PubMed Central PMCID: PMCPMC49875.

43. Bouthenet ML, Souil E, Martres MP, Sokoloff P, Giros B, Schwartz JC. Localization of dopamine D3 receptor mRNA in the rat brain using in situ hybridization histochemistry: comparison with dopamine D2 receptor mRNA. Brain Res. 1991;564(2):203–19. Epub 1991/11/15. doi: 10.1016/0006- 8993(91)91456-b. PubMed PMID: 1839781.

44. Ridray S, Griffon N, Mignon V, Souil E, Carboni S, Diaz J, et al. Coexpression of dopamine D1 and D3 receptors in islands of Calleja and shell of nucleus accumbens of the rat: opposite and synergistic functional interactions. Eur J Neurosci. 1998;10(5):1676–86. Epub 1998/09/29. doi: 10.1046/j.1460- 9568.1998.00173.x. PubMed PMID: 9751140.

45. Sokoloff P, Giros B, Martres MP, Bouthenet ML, Schwartz JC. Molecular cloning and characterization of a novel dopamine receptor (D3) as a target for neuroleptics. Nature. 1990;347(6289):146-51. Epub 1990/09/13. doi: 10.1038/347146a0. PubMed PMID: 1975644.

46. Ryoo HL, Pierrotti D, Joyce JN. Dopamine D3 receptor is decreased and D2 receptor is elevated in the striatum of Parkinson’s disease. Mov Disord. 1998;13(5):788–97. Epub 1998/10/02. doi: 10.1002/mds.870130506. PubMed PMID: 9756147.

47. Quik M, Police S, He L, Di Monte DA, Langston JW. Expression of D(3) receptor messenger RNA and binding sites in monkey striatum and substantia nigra after nigrostriatal degeneration: effect of levodopa treatment. Neuroscience. 2000;98(2):263–73. PubMed PMID: 10854757.

48. Hall H, Halldin C, Dijkstra D, Wikström H, Wise LD, Pugsley TA, et al. Autoradiographic localisation of D3-dopamine receptors in the human brain using the selective D3-dopamine receptor agonist (+)-[3H]PD 128907. Psychopharmacology (Berl). 1996;128(3):240–7. Epub 1996/12/01. doi: 10.1007/s002130050131. PubMed PMID: 8972543.

49. Herroelen L, De Backer JP, Wilczak N, Flamez A, Vauquelin G, De Keyser J. Autoradiographic distribution of D3-type dopamine receptors in human brain using [3H]7-hydroxy-N,N-di-n-propyl-2-aminotetralin. Brain Res. 1994;648(2):222-8. Epub 1994/06/20. doi: 10.1016/0006-8993(94)91121-5. PubMed PMID: 7922537.

50. Landwehrmeyer B, Mengod G, Palacios JM. Dopamine D3 receptor mRNA and binding sites in human brain. Brain Res Mol Brain Res. 1993;18(1- 2):187–92. Epub 1993/04/01. doi: 10.1016/0169-328x(93)90188-u. PubMed PMID: 8097550.

51. Murray AM, Ryoo HL, Gurevich E, Joyce JN. Localization of dopamine D3 receptors to mesolimbic and D2 receptors to mesostriatal regions of human forebrain. Proc Natl Acad Sci U S A. 1994;91(23):11271–5. Epub 1994/11/08. doi: 10.1073/pnas.91.23.11271. PubMed PMID: 7972046; PubMed Central PMCID: PMCPMC45209.

52. Suzuki M, Hurd YL, Sokoloff P, Schwartz JC, Sedvall G. D3 dopamine receptor mRNA is widely expressed in the human brain. Brain Res. 1998;779(1- 2):58–74. Epub 1998/02/25. doi: 10.1016/s0006-8993(97)01078-0. PubMed PMID: 9473588.

53. Piggott MA, Marshall EF, Thomas N, Lloyd S, Court JA, Jaros E, et al. Striatal dopaminergic markers in dementia with Lewy bodies, Alzheimer’s and Parkinson’s diseases: rostrocaudal distribution. Brain. 1999;122 ( Pt 8):1449–68. Epub 1999/08/04. doi: 10.1093/brain/122.8.1449. PubMed PMID: 10430831.

54. Lévesque D, Martres MP, Diaz J, Griffon N, Lammers CH, Sokoloff P, et al. A paradoxical regulation of the dopamine D3 receptor expression suggests the involvement of an anterograde factor from dopamine neurons. Proc Natl Acad Sci U S A. 1995;92(5):1719–23. Epub 1995/02/28. doi: 10.1073/pnas.92.5.1719. PubMed PMID: 7878047; PubMed Central PMCID: PMCPMC42591.

55. Morissette M, Goulet M, Grondin R, Blanchet P, Bédard PJ, Di Paolo T, et al. Associative and limbic regions of monkey striatum express high levels of dopamine D3 receptors: effects of MPTP and dopamine agonist replacement therapies. Eur J Neurosci. 1998;10(8):2565–73. Epub 1998/10/10. doi: 10.1046/j.1460-9568.1998.00264.x. PubMed PMID: 9767387.

56. Visanji NP, Fox SH, Johnston T, Reyes G, Millan MJ, Brotchie JM. Dopamine D3 receptor stimulation underlies the development of L-DOPA-induced dyskinesia in animal models of Parkinson’s disease. Neurobiol Dis. 2009;35(2):184–92. Epub 2009/01/03. doi: 10.1016/j.nbd.2008.11.010. PubMed PMID: 19118628.

57. Payer DE, Guttman M, Kish SJ, Tong J, Adams JR, Rusjan P, et al. D3 dopamine receptor-preferring [11C]PHNO PET imaging in Parkinson patients with dyskinesia. Neurology. 2016;86(3):224–30. Epub 2016/01/01. doi: 10.1212/wnl.0000000000002285. PubMed PMID: 26718579; PubMed Central PMCID: PMCPMC4733157.

58. Boeckler F, Gmeiner P. Dopamine D3 receptor ligands: recent advances in the control of subtype selectivity and intrinsic activity. Biochim Biophys Acta. 2007;1768(4):871–87. Epub 2007/02/06. doi: 10.1016/j.bbamem.2006.12.001. PubMed PMID: 17274946.

59. Newman AH, Beuming T, Banala AK, Donthamsetti P, Pongetti K, LaBounty A, et al. Molecular determinants of selectivity and efficacy at the dopamine D3 receptor. J Med Chem. 2012;55(15):6689–99. Epub 2012/05/29. doi: 10.1021/jm300482h. PubMed PMID: 22632094; PubMed Central PMCID: PMCPMC3415572.

60. Boeckler F, Gmeiner P. The structural evolution of dopamine D3 receptor ligands: structure-activity relationships and selected neuropharmacological aspects. Pharmacol Ther. 2006;112(1):281–333. Epub 2006/08/15. doi: 10.1016/j.pharmthera.2006.04.007. PubMed PMID: 16905195.

61. Heidbreder CA, Newman AH. Current perspectives on selective dopamine D(3) receptor antagonists as pharmacotherapeutics for addictions and related disorders. Ann N Y Acad Sci. 2010;1187:4–34. Epub 2010/03/06. doi: 10.1111/j.1749-6632.2009.05149.x. PubMed PMID: 20201845; PubMed Central PMCID: PMCPMC3148950.

62. Ehrlich K, Götz A, Bollinger S, Tschammer N, Bettinetti L, Härterich S, et al. Dopamine D2, D3, and D4 selective phenylpiperazines as molecular probes to explore the origins of subtype specific receptor binding. J Med Chem. 2009;52(15):4923–35. Epub 2009/07/18. doi: 10.1021/jm900690y. PubMed PMID: 19606869.

63. Zhang A, Neumeyer JL, Baldessarini RJ. Recent progress in development of dopamine receptor subtype-selective agents: potential therapeutics for neurological and psychiatric disorders. Chemical reviews. 2007;107(1):274–302. Epub 2007/01/11. doi: 10.1021/cr050263h. PubMed PMID: 17212477.

64. da Silva-Júnior FP, Braga-Neto P, Sueli Monte F, de Bruin VM. Amantadine reduces the duration of levodopa-induced dyskinesia: a randomized, double-blind, placebo-controlled study. Parkinsonism Relat Disord. 2005;11(7):449–52. Epub 2005/09/13. doi: 10.1016/j.parkreldis.2005.05.008. PubMed PMID: 16154788.

65. Wolf E, Seppi K, Katzenschlager R, Hochschorner G, Ransmayr G, Schwingenschuh P, et al. Long-term antidyskinetic efficacy of amantadine in Parkinson’s disease. Mov Disord. 2010;25(10):1357–63. Epub 2010/03/04. doi: 10.1002/mds.23034. PubMed PMID: 20198649.

66. Luginger E, Wenning GK, Bösch S, Poewe W. Beneficial effects of amantadine on L-dopa-induced dyskinesias in Parkinson’s disease. Mov Disord. 2000;15(5):873–8. Epub 2000/09/29. doi: 10.1002/1531- 8257(200009)15:5<873::aid-mds1017>3.0.co;2-i. PubMed PMID: 11009193.

67. Fox SH, Katzenschlager R, Lim SY, Ravina B, Seppi K, Coelho M, et al. The Movement Disorder Society Evidence-Based Medicine Review Update: Treatments for the motor symptoms of Parkinson’s disease. Mov Disord. 2011;26 Suppl 3:S2–41. Epub 2011/11/02. doi: 10.1002/mds.23829. PubMed PMID: 22021173.

68. Perez-Lloret S, Rascol O. Efficacy and safety of amantadine for the treatment of L-DOPA-induced dyskinesia. Journal of neural transmission (Vienna, Austria : 1996). 2018;125(8):1237–50. Epub 2018/03/08. doi: 10.1007/s00702-018-1869-1. PubMed PMID: 29511826.

69. Tanner CM, Pahwa R, Hauser RA, Oertel WH, Isaacson SH, Jankovic J, et al. EASE LID 2: A 2-Year Open-Label Trial of Gocovri (Amantadine) Extended Release for Dyskinesia in Parkinson’s Disease. J Parkinsons Dis. 2020;10(2):543–58. Epub 2020/01/14. doi: 10.3233/jpd-191841. PubMed PMID: 31929122; PubMed Central PMCID: PMCPMC7242830.

